# A subset of mouse hippocampus CA1 pyramidal neurons learns sparse synaptic input patterns

**DOI:** 10.1101/2024.12.05.626959

**Authors:** Anzal K Shahul, Upinder Singh Bhalla

## Abstract

Synaptic plasticity in the hippocampus is fundamental to learning and memory, yet few studies examine how pattern learning occurs across multiple synapses. Such cross-synapse learning is fundamental to emergent properties of pattern discrimination and generalisation, which depend on assumptions about independence of plasticity and linearity of summation. We used sparse optogenetic spatio-temporal ‘pattern stimulation in the CA3 coupled with postsynaptic depolarization to elicit plasticity on CA1 pyramidal neurons, and found that ‘trained’ patterns were selectively strengthened, but only in a subset of postsynaptic cells. Increased resting membrane potential and background mini-EPSP rates were predictive of learner cells. Summation following plasticity became more linear in learners compared to non-learners, consistent with the observed elevated post-stimulus hyperpolarization on non-learner cells. Thus our exploration of biologically plausible sparse activity supports pattern-selective learning, but in a heterogeneous manner modulated by both cell-intrinsic and network features.

## Introduction

### Sparse activation of the hippocampal CA3-CA1 circuit

By combining sensory, emotional, and contextual components into coherent memories, the hippocampus plays a crucial role in the encoding, retrieval, and consolidation of events (Eichenbaum, 2017; Knierim, 2015). Place cells in the hippocampal CA1 are the prototypical example of convergence of cues through sparse activation in the CA3, leading to CA1 pyramidal neuron (PN) plasticity and functions of spatial memory and navigation (Dong et al., 2021).

Long-term potentiation (LTP) at synapses in the hippocampus CA3-CA1 circuit is believed to be a major component of network learning, and many stimulation modalities can induce LTP (Bittner et al., 2017). Common approaches are field electrode stimulation, glutamate uncaging and wide-field optogenetic stimulation (Bortolotto et al., 2001; Jain et al., 2024).

Less commonly used optogenetic techniques include 2-photon stimulation and patterned illumination using DMDs or SLMs (Bègue et al., 2013; Hochbaum et al., 2014).

The classical approach for induction of slice LTP is to stimulate the Shaffer collaterals using large electrodes (Ajay and Bhalla, 2004). This precisely controls timing, but recruits an unphysiologically large number of axons mostly from the CA3, and a less well defined population of inhibitory-neuron axons.

Glutamate uncaging, in contrast, achieves synapse-level specificity using photolysis to release caged glutamate at specific synaptic locations (Jain et al., 2024). This bypasses presynaptic activity, thus restricting plasticity mechanisms and modulation to postsynaptic events. Further, glutamate uncaging is exclusive to the targeted synapse, thus does not recruit the population of interneurons which provide inhibitory balance.

Patterned optogenetic stimulation avoids many of these drawbacks, by combining target-specificity and spatial pattern selectivity, while also retaining temporal precision. One can selectively express opsins in CA3 pyramidal neurons and choose spatial and temporal patterns to match physiological levels of activity (Oishi et al., 2019). We adopt this approach using DMD-based patterned illumination, to preserve E-I balance and also to facilitate fine-grain analysis of network phenomena such as multi-input summation, pattern discrimination and generalisation.

### Heterogeneity in the CA1 PN population

CA1 pyramidal neurons exhibit notable variation in their intrinsic membrane characteristics, morphology, and synaptic connections. There are large-scale graded changes across the dorsal-ventral axis (Milior et al., 2016), but our current study homes in on finer-grained heterogeneity within the dorsal CA1. One clear level of heterogeneity is seen by way of molecular markers, where there is evidence for two distinct populations of CA1 neurons depending on expression of CaMKII-alpha (Sarkar et al., 2021). At the physiological level, several collections of CA1 PN recordings exhibit a diversity of morphological and physiological features (Anal Kumar et al., 2024; Basak and Narayanan, 2020; Milior et al., 2016; Romani et al., 2024). These include differences in passive properties, firing patterns and excitability, which affect their responses to synaptic inputs. At a still finer-grain level, single-synapse recordings using glutamate uncaging support this picture of a range of baseline and plasticity profiles for synapses (Weber et al., 2016).

Despite this heterogeneity at the single-cell and synapse level, CA1 PNs also exhibit remarkably precise E-I balance (Bhatia et al., 2019), suggesting that function is closely maintained in the midst of degeneracy (Hutt et al., 2023). It is therefore interesting to see how EI balance and summation evolve over the course of network plasticity, and whether the plasticity itself is heterogeneous.

In our study we used optogenetics to deliver sparse spatially patterned stimulation to CA3 PNs to explore the ability of postsynaptic CA1 PNs to perform pattern separation and pattern completion following induced synaptic plasticity. Plasticity was diverse: about half of the recorded neurons exhibited pattern-specific potentiation, yet there was little divergence from EI balance and continued divisive normalisation in all cells. Multi-input summation remained sublinear following plasticity.

## Results

In the experiments below, we first characterise the optical input-output properties of our optogenetically stimulated slice preparation. We ‘train’ the CA1 PN on one of three spatial optical patterns used to stimulate the CA3 PN layer. We measure the range of plasticity, show it is selective to spatial input, and relate it to cellular physiological properties. Finally, we break down the plasticity changes to individual points in the spatial pattern and show how summation changes during pattern learning.

### Optical stimulus patterns at CA3 PNs elicit reliable EPSPs on CA1 PN soma

We delivered spatiotemporally patterned stimuli using an LED micromirror device with 0.49 *µ*m resolution to stimulate channelrhodopsin-2 (ChR2) expressed using a CA3 pyramidal neuron-specific promoter (Figure 1A-C). We recorded extracellular local field potentials (LFPs) in the CA3 to examine the collective neuronal activity elicited by optogenetic stimulation (Figure 1Di-iii). Simultaneously, we obtained whole-cell patch-clamp recordings from CA1 pyramidal neurons to directly measure the summed EPSPs produced by synaptic inputs from the stimulated CA3 region and the associated feedforward inhibitory responses (Figure 1Ci-iii). We observed a range of EPSP amplitudes and latencies(Figure 1Ei, Fi). Notably, the LFP trial-to-trial variability was much smaller than that of EPSP, suggesting that the stimulus amplitude was consistent and hence only a small contribution to EPSP amplitude differences (Figure 1 G). We also confirmed that the latency for LFP was short (∼4 ms) and tight (σ=2.66 ms) (Fig Fii), extending the uniformity of stimuli to the time domain. Thus the optical stimulus method was reliable and precise in activating the CA3 and delivering patterned stimuli to downstream neurons.

**Figure 1.**
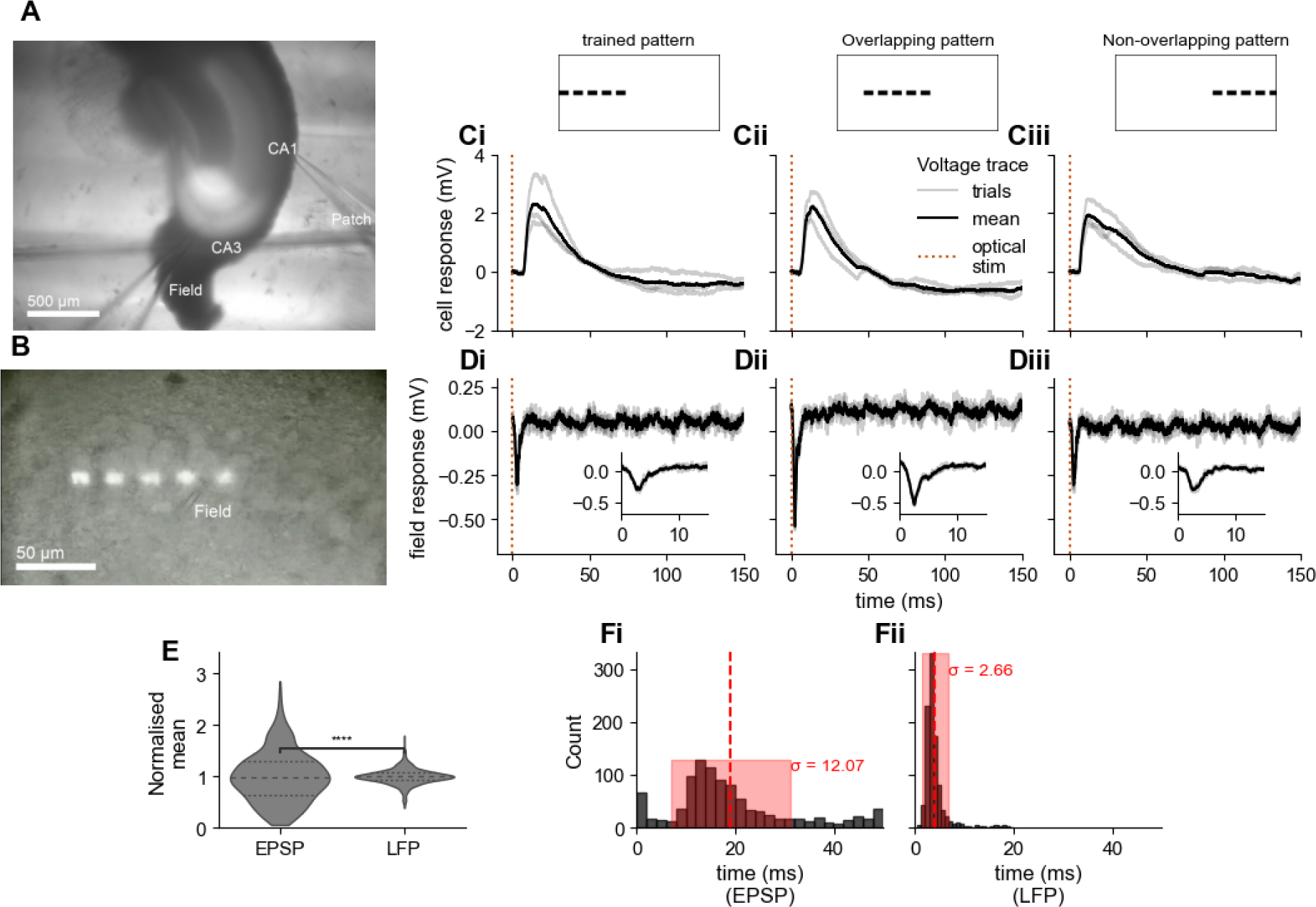
Characterization of hippocampal CA1 responses to patterned optogenetic stimulation. **(A)** Brightfield (DIC + fluorescence) image of a hippocampal slice showing the anatomical regions CA1 and CA3, the location of the patch-clamp recording electrode (Patch), and the field recording site (Field) **(B)** DIC brightfield image of spatial light stimulation pattern in the CA3 region. LFP electrode location is indicated as *Field*, and the visible CA3 PN soma (contrast adjusted for better visibility). **(Ci-Ciii)** CA1 PN EPSPs in response to optogenetic stimulation of trained, overlapping, and non-overlapping input patterns respectively. Individual trial traces are shown in gray, and the trial-averaged response is shown in black. The dotted orange vertical line indicates the onset of optical stimulation. **(Di-Diii)** Field potential responses (mV) to optogenetic stimulation of trained, overlapping, and non-overlapping input patterns respectively. Individual trial traces are shown in gray, and the trial-averaged response is shown in black. The inset shows a magnified view of the initial 10 ms response. The dotted orange vertical line marks the onset of optical stimulation. **(E)** Violin plots comparing the variance in normalized EPSP and LFP responses. LFPs are much tighter. Statistical significance is indicated (**** p < 0.0001, with Levene’s test). **(F)** Optically stimulated LFP timings are precise and rapid compared to EPSPs (**** p < 0.0001, with Levene’s test). The red dashed line indicates the mean, and the red shading represents the standard deviation (σ in ms).

### Optically induced CA1 LTP lies on a continuum

We next investigated plasticity triggered by our spatially controlled, sparse stimulation protocol. We used three spatial patterns arranged with light spots along a line, such that the first two patterns had some overlap but the first and third were well-separated (Figure 2 B). The first pattern was used for plasticity induction, the second (overlapping) pattern for testing pattern completion, and the third, well separated pattern to test for changes at non-induced synapses. Our training protocol is indicated in Figure 2A and illustrative responses in 2C. Briefly, we obtained baseline EPSP responses, induced plasticity by pairing presynaptic activation with postsynaptic depolarization in an STDP-like protocol with 100 pairings repeated five times, and then monitored EPSPs to the test patterns over at least 30 minutes following LTP induction.

**Figure 2.**
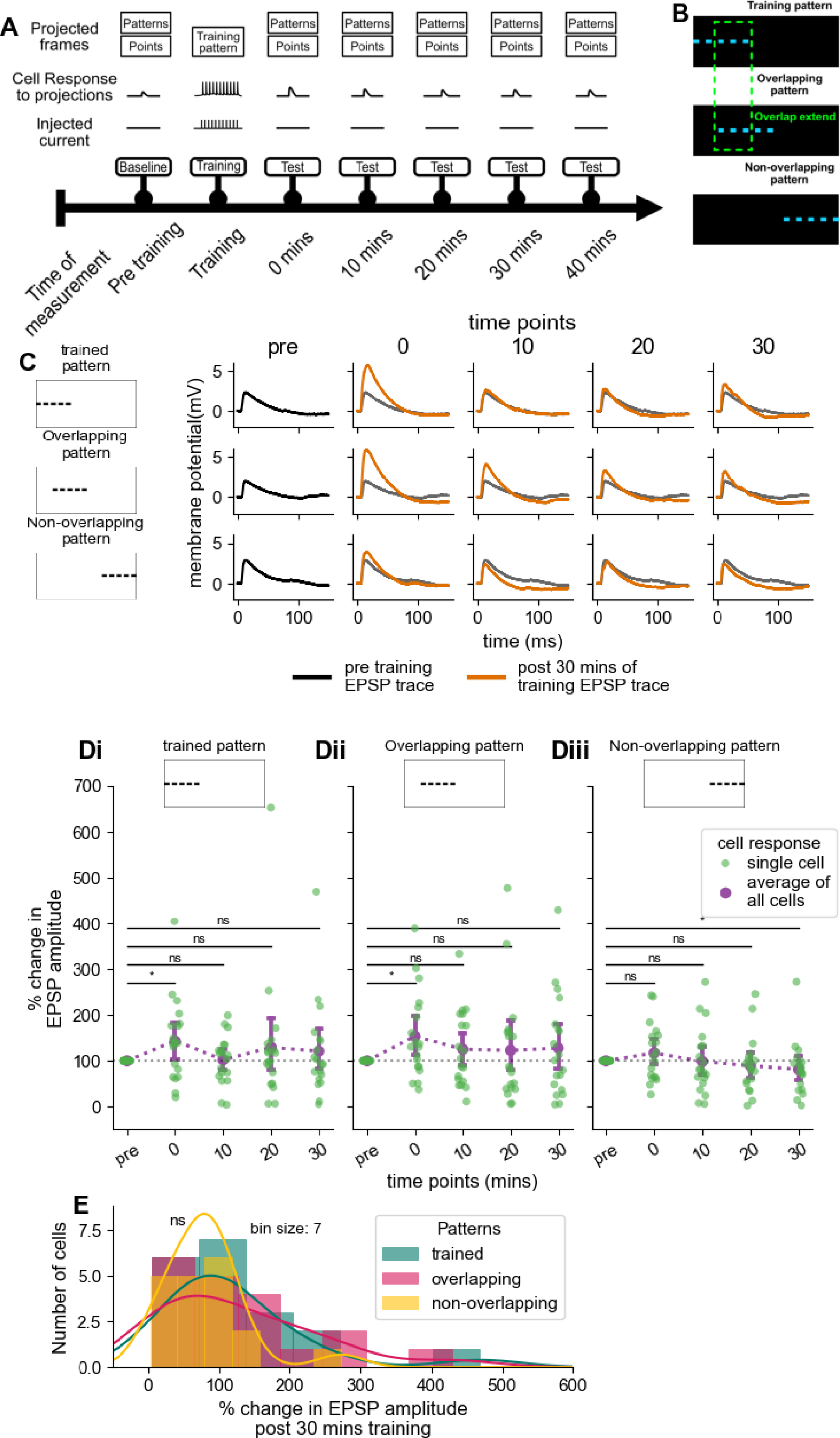
Optogenetically induced plasticity in CA1 is heterogeneous. **(A)** Schematic timeline of the experimental protocol, showing the different phases of measurement at multiple time points (0, 10, 20, and 30 minutes). Top row: stimulus patterns. Second row: intracellular potentials. Third row: injected current. Bottom row with arrow: phase of training. LTP was induced by pairing of the training pattern with postsynaptic depolarization. EPSPs were recorded for all three patterns before and after training. **(B)** Schematic illustrating the three categories of patterns used for stimulation: trained, overlapping, and non-overlapping patterns. The cyan bright spots indicate the illuminated regions of size 13 x 7 *µ*m under 40x objective. The extent of overlap is indicated in the green dotted box. **(C)** Representative traces of membrane potential responses to three stimulation patterns: trained, overlapping, and non-overlapping at pre-training (left, black), and at multiple time points following training (0, 10, 20, and 30 minutes, orange). All traces have baseline resting membrane potential (RMP) subtracted, and each trace represents an average of three trials. **(Di-Diii)** Changes in EPSP amplitude in response to the trained (Di), overlapping (Dii), and non-overlapping (Diii) patterns(n=18). Each green dot represents a single cell’s response, averaged over three trials. Baselines are normalised to 100%. The purple line and error bars represent the mean EPSP amplitude ± SEM across cells. Most distributions do not differ significantly from baseline as measured by the Wilcoxon signed-rank test, (* = p<0.05, ns = non-significant). **(D)** Distribution of EPSP amplitudes 30 minutes post-training, normalized to baseline. The peaks are not significantly different (pairwise KS test).

Contrary to our expectations, CA1 PN responses following plasticity were highly heterogeneous. Eight neurons showed LTP, with a sustained increase in EPSP amplitude 30 minutes after training, whereas 12 neurons did not show clear plasticity or even a reduction in EPSP (Figure 2 D,E). Thus our sparse stimulation protocol yields a spectrum of plasticity responses, from potentiation to depression, in contrast to established Schaffer collateral stimulus protocols (Abraham and Huggett, 1997; Ouyang et al., 1999). The non-overlapping control pattern did not increase EPSP amplitude over 30 minutes, but instead reduced in amplitude in 12 out of 18 cells (Figure 2Diii, E), indicating that synaptic strength increases were limited to stimulus-paired synapses during training (p<0.05, Wilcoxon signed rank test). In the same recordings, we also evaluated pattern-selective plasticity at a series of timepoints after induction and before the 30 minute LTP mark. Immediately after plasticity induction (0 minutes time point, as shown in Figure 2 A), all neurons had elevated EPSP amplitudes in response to trained as well as overlapping patterns (p=0.045 and p=0.045, respectively, Wilcoxon signed rank test). In contrast, the non-overlapping patterns did not elicit a response greater than baseline. Subsequent test readouts did not exhibit significant potentiation when averaged over the entire dataset (Figure 2 Di, Dii and Diii, ns, with Wilcoxon signed rank test). Instead, there was considerable scatter in the readouts for all patterns

In summary, we were able to elicit pattern-selective plasticity which was consistent across cells at short time points but became heterogenous at the 30-minute time-scale of LTP.

### CA1 pyramidal cell learning is correlated with resting membrane potential and mEPSP frequency

We next examined a range of physiological features of the CA1 PNs to see if they were correlated with LTP outcomes. First, we categorised ‘learners’ as cells which retained a significantly higher EPSP over baseline at the 30 minute time-point after LTP induction (Fig 3A, B). By this criterion, there were 8 learners and 10 non-learners. Other criteria for LTP, such as reduction in the EPSP onset time and area under the curve and area under curve, gave the same categorization of learners and non-learners (Supplementary Figure 1A, G). We found noticeable EPSP waveform differences between learners and non-learners. In particular, non-learners showed a characteristic post-stimulus hyperpolarization after the initial EPSP (Fig 3C).

**Figure 3.**
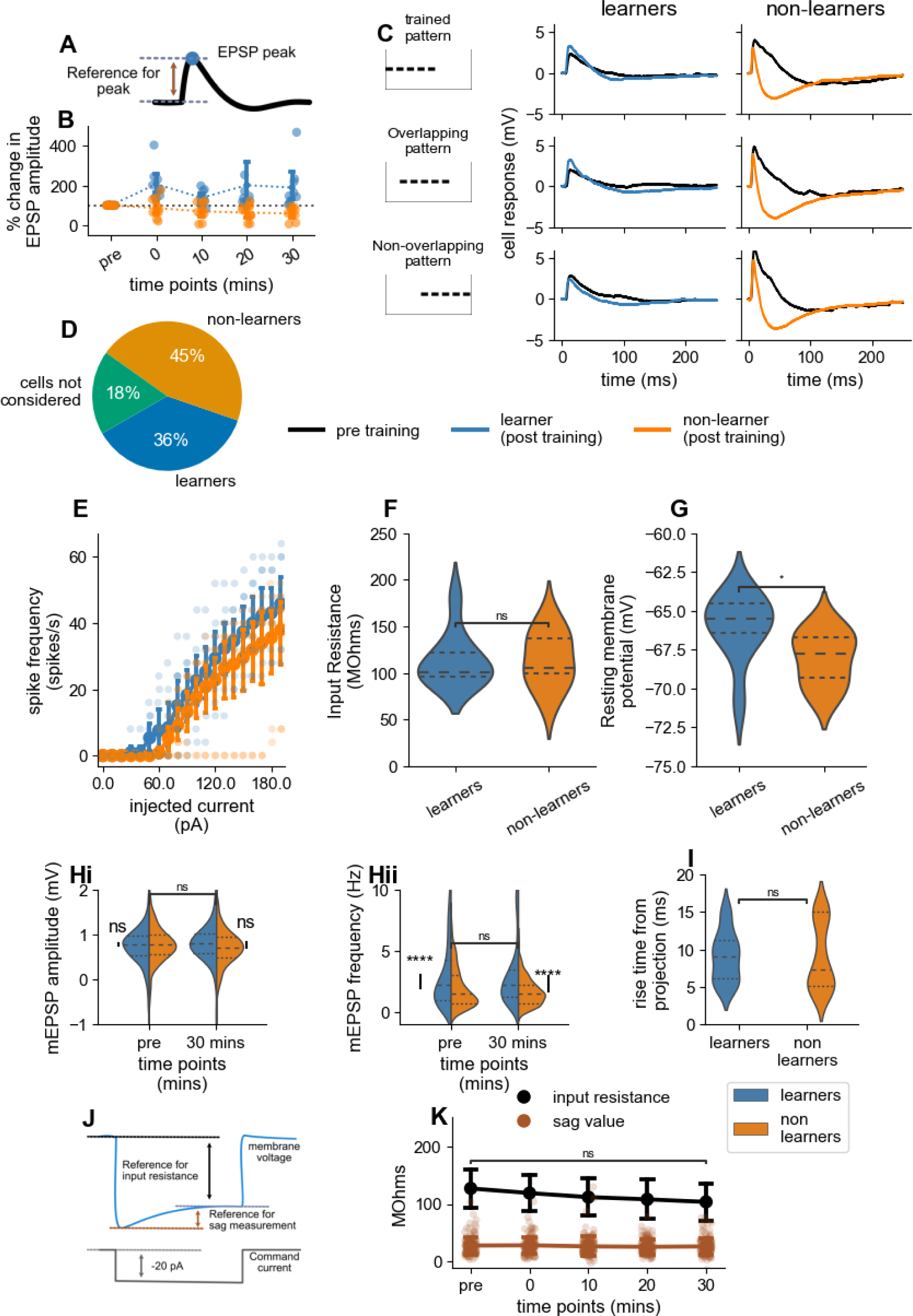
Changes in synaptic responses and intrinsic properties of CA1 neurons during pattern learning. All comparisons shown in the figure are made at 30 minutes post-training unless otherwise specified. **(A)** Schematic of measurement of EPSP peak as a measure of synaptic response. **(B)** Scatter plot showing the percentage change in EPSP amplitude over time for learners (blue) and non-learners (orange) at different time points (pre, 0, 10, 20, and 30 minutes). Error bars represent mean ± SEM. **(C)** Representation of the trained, overlapping, and non-overlapping patterns. The right panels show representative membrane potential responses for learners (left) and non-learners (right). Black traces represent baseline responses (pre-training), while orange traces represent post-training responses. **(D)** Pie chart summarizing the distribution of cells classified as learners (36%, n=8), and non-learners (45%, n=10). Four cells (18%) were excluded because their EPSP amplitudes were less than 0.5 mV, or because they spiked after training. **(E)** Input-output relationship between spike frequency and injected current for learners (blue) and non-learners (orange). The curves did not differ. Error bars represent mean ± SEM. **(F)** Comparison of input resistance (MΩ) between learners and non-learners. No significant differences were observed (Mann-Whitney U test). **(G)** Comparison of resting membrane potential between learners and non-learners. Learners exhibited a significantly more depolarized resting membrane potential compared to non-learners (p = 0.025,Mann-Whitney U test). **(H)** Miniature EPSP (mEPSP) analysis in learners and non-learners, compared before and 30 minutes after training.**(Hi)** mEPSP amplitude comparison shows no significant differences. **(Hii)** Comparison of mEPSP frequency shows a significant difference between the learners and non learners (p<0.0001, Mann-Whitney U test). The pre and post-training frequency do not differ significantly. **(I)**. Response rise time, measured as time from delivery of optical stimulus till reaching 0.25 mV depolarization. Learners and non-learners did not differ (Mann-Whitney U test). **(J)** Schematic of the electrophysiological protocol used to measure input resistance and sag potential. **(K)** Input resistance (black) and sag potential (brown) measured over time relative to baseline (pre-training). Data are presented as mean ± SEM. No significant changes were observed over the duration of training (Wilcoxon signed rank test).

We evaluated firing properties, input resistance, and resting membrane potentials to see if intrinsic cellular properties differed between learners and non-learners (Fig 3E-J). No significant variations in firing rates or input resistance were found between the two groups, suggesting that intrinsic excitability does not impact neuron learning (ns with Mann-Whitney U Test). However, learners had a more positive resting potential than non-learners (Fig 3G, p=0.025 with Mann-Whitney U Test). The sag response, a measure of intrinsic membrane plasticity related to hyperpolarization-activated HCN channels, also did not change after plasticity induction (Figure 3 K, (ns, Wilcoxon signed rank test)).

To assess synaptic connectivity contributions to learning, we next examined mini EPSPs (mEPSPs) of non-stimulus periods before and after LTP induction. To avoid the tail of EPSPs following synaptic input, we considered the membrane potential measured one second after the optical stimulation at CA3. Before training, there was no significant difference in mEPSP amplitude between learners and non-learners, but there was a significant difference (p<0.0001) in mEPSP frequency. At the 30 minutes time point following plasticity induction, we saw a significant difference in mEPSP amplitude between learners and non-learners (p=0.0005 with Mann-Whitney U test), and also a significant difference between the mEPSP frequency of learners and non-learners(p<0.0001 and p<0.00001 with Mann-Whitney U test)). The difference in mEPSP frequency even before learning suggests a potential connectivity difference from the presynaptic terminals to CA1 PNs for the learners and non-learners.

Membrane voltage dynamics did not differ between learners and non-learners during training, as measured by the rise time for membrane potential to 0.25 mV after optical stimulus (Fig 3 I).

We also observed potentiation in LFP recorded in the CA3 (Supplementary Figure 2). The CA3 LFP has an indirect relationship with number of active CA3 axons, since the placement of the electrode also picks up dendritic and synaptic currents, and further there is a nonlinear relationship between LFP and number of active fibres (Lindén et al., 2010). We asked if the outcome of the analysis would change in the most restrictive case, that is, if the number of active Schaffer collateral axons was proportional to LFP (Supplementary Figure 2). While the number of learners was reduced from eight to six, none of the key conclusions about plasticity, pattern selectivity or summation change (Supplementary Figure 3 and 4).

Overall, CA1 PNs can be categorized as learners or non-learners. These groups have similar intrinsic properties, but there are two significant differences that correlate with subsequent learning. First, learners have a higher membrane potential. Second, higher mEPSP frequency strongly correlates with learning.

### Learner neurons exhibit pattern selectivity contributed by multiple synapses

We next examined the degree to which neurons were selective for spatially overlapping input patterns. We designed our three input patterns as linear arrays located over the CA3 cell body layers, to minimise activation of dendrites and axonal bundles, which would have lessened the spatial specificity of the stimulus patterns (methods). In brief, for learner cells both the stimulus and overlap patterns potentiated, while the non-overlap pattern did not (Figure 4 A, C). This suggests that learners’ LTP plasticity was unique to synaptic inputs directly involved in training. Unexpectedly, the overlap patterns underwent as much plasticity as the trained patterns. As considered in the discussion, this could be due to stimulus overlap. For non-learner cells there was no pattern selectivity. Instead, all patterns led to depression (Figure 4 B, C).

**Figure 4.**
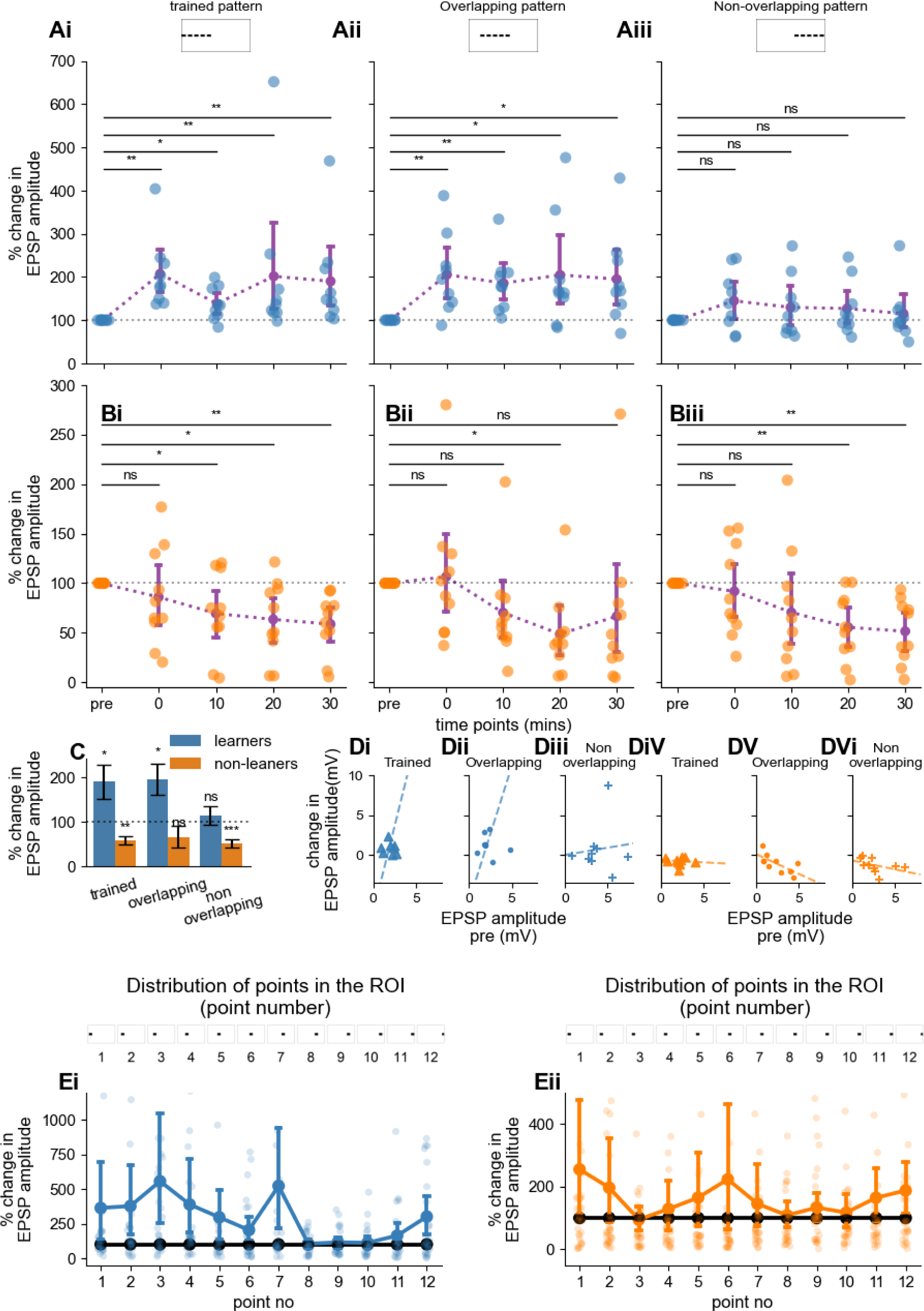
EPSP amplitude changes following optogenetic stimulation patterns for learners and non-learners. **(Ai-Aiii)** Quantification of change in EPSP amplitude in learner cells (n=8). Amplitudes reported as percentage change normalized to baseline for trained (Ai), overlapping (Aii), and non-overlapping (Aiii) stimulation patterns across time points. Individual points represent single cells, lines and error bars indicate the mean ± SEM. Trained ( p=0.0039) and overlap (p=0.011) patterns have a significant increase by the Wilcoxon signed rank test. **(Bi-Biii)** Same as A for non-learners (orange, n=10). Trained (p=0.0019) and non-overlap (p=0.0019) patterns show a significant decrease, Wilcoxon signed rank test. **(C)** Population summary for A and B, measured at the 30 minute time point.We used the two-tailed t-test to compare the groups with the baseline. For learners, both trained and overlapping patterns show a significant increase(p=0.045 and p=0.026 respectively) while non-overlapping didn’t show any difference.For non learners, both trained and non-overlapping patterns showed a decrease in the EPSP amplitude(p=0.00199 and p=0.00096 respectively) **(D)** Scatter plot and linear regression fit for EPSP amplitude at 30 minutes post-training versus baseline (pre-training) amplitude for individual cells. **(Di-iii)** Learners did not show any correlation with the initial EPSP amplitude to change in EPSP amplitude post training for any patterns. r=0.18, 0.17 and 0.08 with Spearman correlation test, for trained, overlapping and non-overlapping patterns respectively. **(Div-vi)** Non-learners show a negative correlation for the change in amplitude with the initial EPSP amplitude(r=0.08, −0.75 and −0.77 and p=0.829, 0.013 and 0.009 with Spearman correlation test for trained, overlapping and non-overlapping patterns respectively). **(E)** Plasticity for individual points in the spatial patterns. Points 1-5 underwent the LTP stimulus protocol, the overlap pattern was points 3-7, and non-overlap pattern was points 8-12. Error bars represent the mean ± SEM of EPSP. The baseline is normalized to 100% in all plots. **(Ei)** learners. Despite group percent change being higher for trained pattern (points 1-5) and learned pattern (points 3-7), none of the individual points elicited a significantly higher EPSP (Wilcoxon signed rank test). **(Eii)** non-learners. Neither the individual points nor the groupings into patterns were significantly different from baseline EPSP.

In order to test whether initial synaptic strength influenced plasticity, we compared each cell’s initial EPSP with the change in EPSP amplitude post-training. Initial synaptic weight did not correlate with potentiation or depression levels for learners (ns with spearman correlation). For the non-learners, the change in EPSP amplitude negatively correlated with the initial synaptic weight for non-overlapping patterns (r=-0.77, p=0.009 with spearman correlation). In summary, learner neurons exhibited spatial pattern selectivity in their learning, and this capacity was independent of the initial summed synaptic strength.

### All patterns elicit increased post-stimulus hyperpolarisation(PSH) following training

Post-stimulus hyperpolarisation (PSH) increased across all synaptic input patterns after training, both for learners and non-learners (Wilcoxon signed rank test) (Figure 5A,B and C). The one exception was for the trained pattern in non-learners. The PSH built up after the immediate post-stimulus period, and was sustained for the entire duration of the recording (Figure 5, B, C, D). The final PSH did not depend on the initial EPSP amplitude (Figure 5 E). We further checked if there was spatial specificity of PSH in relation to the stimulus points. Surprisingly, the rise in PSH was not restricted to individual points in the input patterns, either for learners or non-learners (Wilcoxon signed rank test) (Figure 5F).

**Figure 5.**
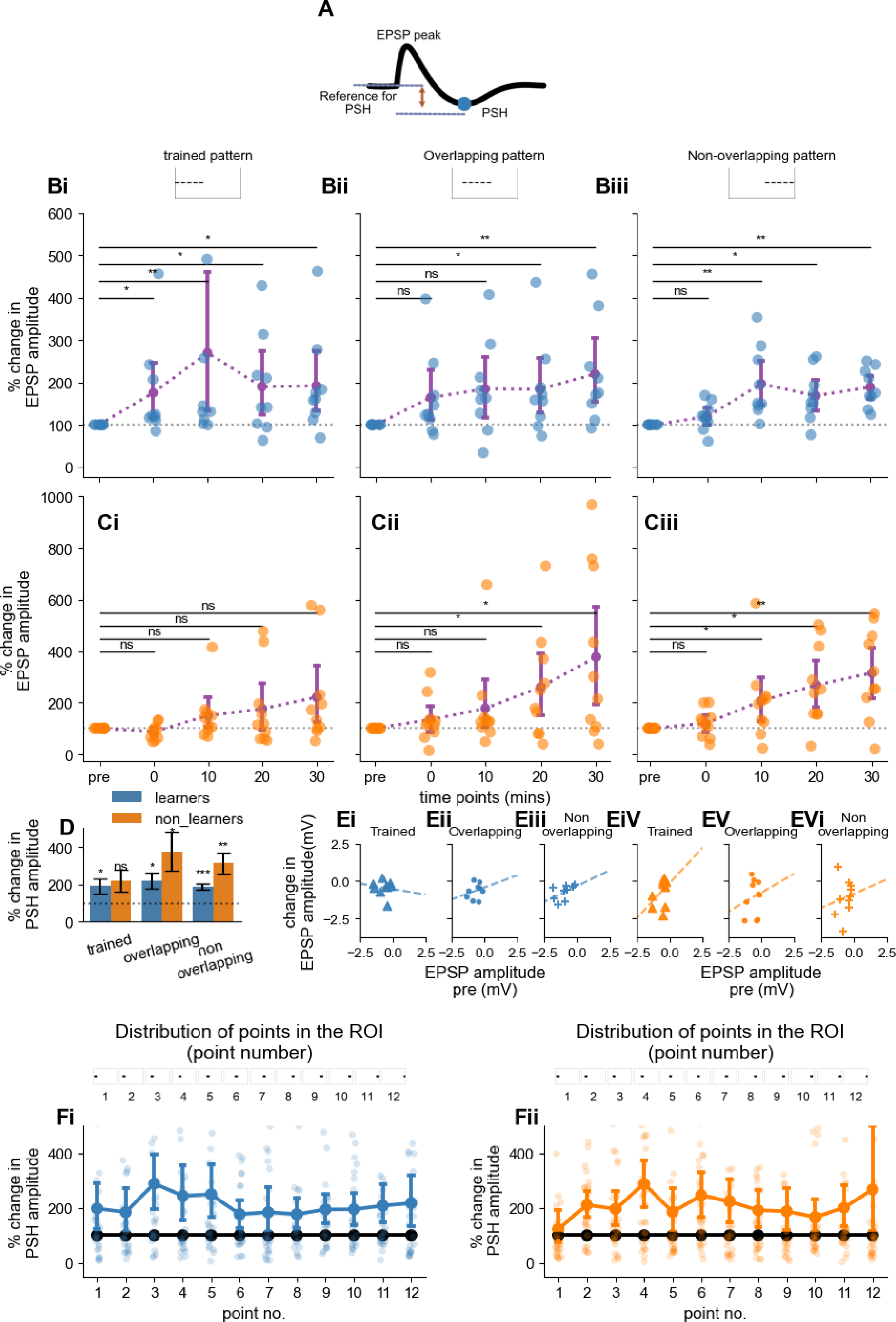
Post-stimulus hyperpolarization (PSH) changes. **(A)** Schematic representation of the EPSP waveform showing the peak and post-stimulus hyperpolarization (PSH), used to quantify changes following stimulation. **(B)** Quantification of the percentage change from baseline in PSH amplitude for the trained **(Bi)**, overlapping **(Bii)**, and non-overlapping **(Biii)** stimulation patterns across time points (pre-training, 0, 10, 20, and 30 minutes post-training) in learner cells (blue, n=8). Each point represents an individual cell, and lines with error bars indicate the mean ± SEM. Trained (p=0.0019), overlap (p=0.0078) and non-overlap (p=0.004) patterns show a significant increase (Wilcoxon signed rank test). The purple dashed lines represent the trend of PSH amplitude changes over time. **(C)** Similar to B, for non-learners.overlapping (p= 0.027) and non-overlap (p=0.0058) patterns show a significant decrease, Wilcoxon signed rank test. **(D)** Bar plot summarizing the average percentage change in PSH for learners (blue) and non-learners (orange) in response to the trained, overlapping, and non-overlapping patterns. Error bars represent the mean ± SEM. The dotted line indicates the baseline response. We see a significant increase across patterns for learners and non-learners except non-learners for the trained pattern (trained pattern: learners; p =0.045, non-learners; p =ns, overlapping pattern: learners; p =0.0018, non-learners; p =0.026, non-overlapping pattern: learners; p =0.000, non-learners; p =0.003, with with two tailed t-tests). **(E)** Scatter plot showing the PSH amplitude at 30 minutes post-training versus baseline (pre-training) for individual cells. Each point represents a cell’s response to a specific stimulation pattern, with data points color-coded by learner status (learners vs. non-learners). Learners showed a negative correlation with the initial EPSP amplitude to change in PSH amplitude post training for trained and non-overlapping (Eiii) but not for overlapping (Eii) patterns (r=-0.73, −0.33 and −0.83, p=0.025, 0.038, 0.005 with Spearman correlation test, for trained(Ei), overlapping (Eii) and non-overlapping (Eiii) patterns respectively). **(Eiv-vi)** Non-learners showed a slight positive correlation with change in PSH to the initial EPSP amplitude for the trained pattern (r=0.26, 0.13 and −0.07 and p=0.467, 0.726 and 0.856 with Spearman correlation test for trained (Eiv), overlapping (Ev) and non-overlapping (Evi) patterns respectively). **(F)** Temporal dynamics of the percentage change in PSH across 12 stimulation points within each pattern for learners (Fi, blue) and non-learners (Fii, orange). Each point corresponds to a specific region of illumination (ROI) in the stimulation pattern, and error bars represent the mean ± SEM. Baseline is normalized to 100%. We did not find any significant difference between the baseline and the EPSP amplitudes at 30 mins time point(Wilcoxon signed rank test).

In summary, the training stimulus elicited an increase in PSH irrespective of learning status, prior PSH for that stimulus, or input pattern, except for the trained pattern in non-learners.

### Plasticity does not alter summation either in learners or non-learners

We next utilised our patterned optical stimulus protocol pre- and post-learning to examine how neuronal summation changes over the course of plasticity. Each of our patterned stimuli was composed of five points of illumination, whose responses we measured individually and in patterns. We analysed the summation in terms of linearity and match to divisive normalisation (Bhatia et al., 2019). In brief, we computed expected response as a linear sum of the response to individual stimulus points in the pattern (Fig 6A). We compared this with the observed response, and used the general normalization equation from (Bhatia et al., 2019)

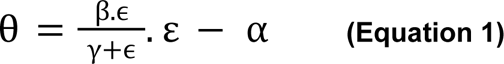

**Figure 6.**
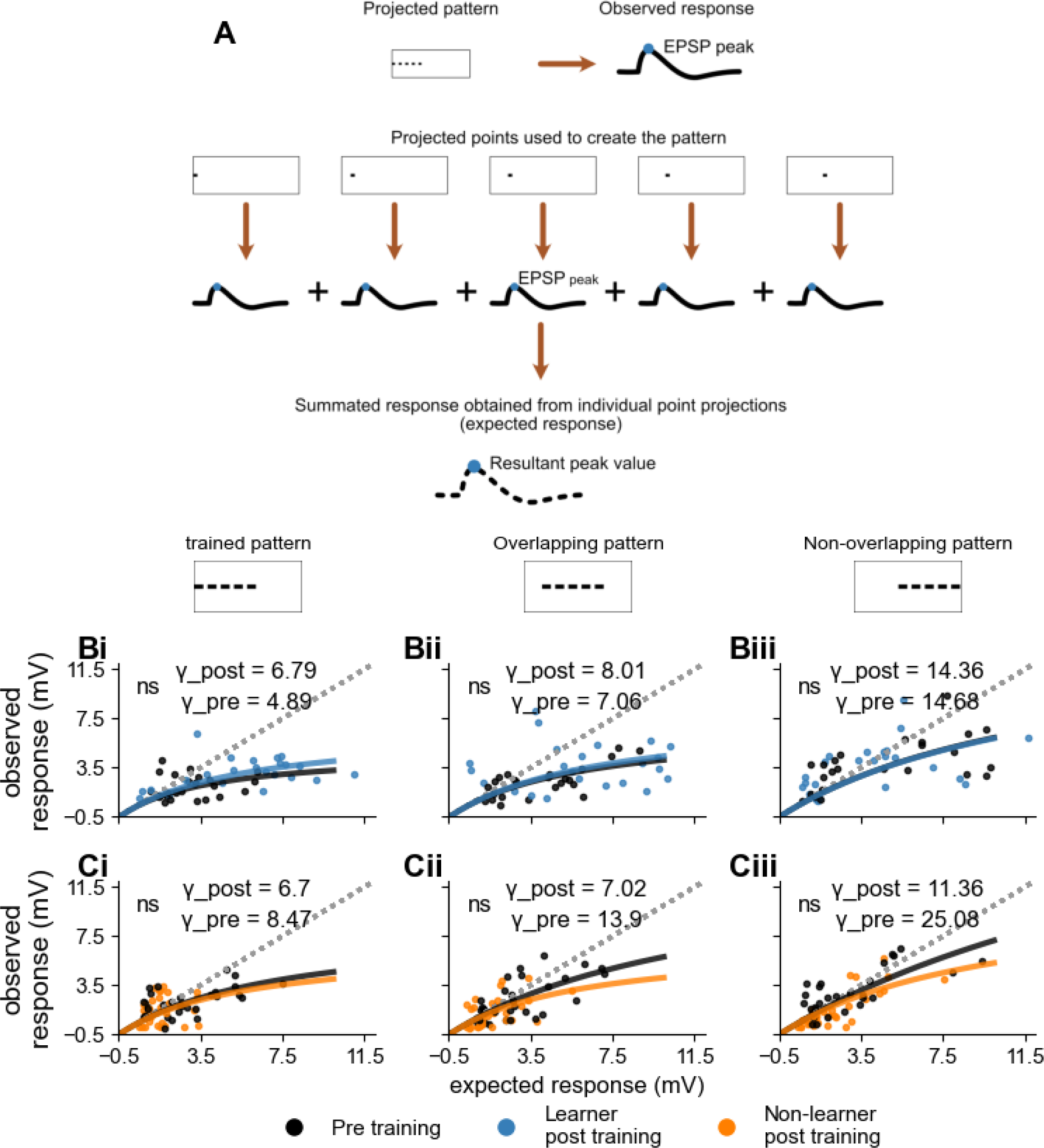
Sublinear Summation and Divisive Normalization in CA1 Neurons Across Different Patterns. **(A)** Schematic illustrating how the expected EPSP peak is obtained by summing the EPSP peak obtained with individual point projections. The resultant distribution of observed vs expected EPSPs is fit to Equation 2 to obtain *γ*. Higher values of *γ* indicate greater linearity. Bootstrap analysis was used to compare γ-pre with γ-post, with the null hypothesis that the pre and post-training distributions were identical (methods). **(B)** Results from learners(n=8, 3 trials per pattern), with fits for γ-pre (black) and γ-post (blue). A linear sum is represented by the dotted line. **(Bi)**For trained pattern (γ-pre = 4.89, γ-post = 6.79, p=ns) **(Bii)** Overlapping pattern (γ-pre = 7.06, γ-post = 8.01, p=ns) and **(Biii)** Non-overlapping pattern (γ-pre = 14.36, γ-post =14.68, p=ns) have no significant change in the summation. **(C)** Results from non-learners (n=10, 3 trials per pattern).**(Ci)** Trained pattern (γ-pre = 6.7, γ-post = 8.47, p=ns). **(Cii)** Overlapping pattern (γ-pre = 7.02, γ-post = 13.9, p=ns) and **(Ciii)** non-overlapping pattern (γ-pre = 11.36, γ-post = 25.08,p=ns) γ does not change significantly. We also did not see any significant difference between the gammas of trained and non-overlapping patterns.

where:

- **θ** is the observed response,
- **ε** is the expected response (sum of responses to individual point projections),
- **α** is the subtractive inhibition parameter,
- **β** is the divisive inhibition parameter,
- **γ** is the normalization parameter.

For Divisive Normalization, **α = 0**, **β = 1**, and **γ** is treated as a free parameter. Thus we obtain the simplified equation:

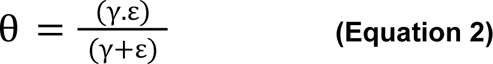

We used Equation 2 to fit the points before and after LTP (Figure 6 B, C). As expected from previous work (Bhatia et al., 2019) we found that divisive normalization gave a good fit to the observations for all the plots (Figure 6 B). We parameterized summation linearity measured by the γ parameter, and compared it before and after plasticity induction using the bootstrap method (Figure 6B). Notably, none of the γ terms for learners or non-learners, nor for the different patterns, showed a significant difference (Figure 6 B,C). However, in the LFP-scaled dataset the trained pattern for learners became more linear (Supplementary Figure 4).

In summary, the analysis of summation showed that all pattern responses in all neurons obeyed divisive normalization before and after the LTP stimulus, notwithstanding the diversity of plasticity profiles.

## Discussion

We investigated synaptic plasticity and pattern learning in mouse hippocampal CA1 PNs using sparse spatially patterned optogenetic stimulation of CA3 PNs. We find that CA1 neurons are heterogeneous: “learners” demonstrated long-term potentiation and developed pattern discrimination, but “non-learners” did not. This categorization of CA1 PNs was predicted by correlations of learners with elevated resting membrane potential and mEPSP frequency. There was uniform post-stimulus hyperpolarization across all patterns following training, which suggests this is a mechanism for balancing excitability. Despite the heterogeneity in plasticity profiles, all cells sum sub-linearly before and after plasticity, and plasticity does not alter the degree of sub-linearity..

### A synapse-specific basis for pattern learning

We observed learning for trained and overlap patterns in some cells, but in the same cells non-overlap patterns did not potentiate (Figures 2, 3, 4). This is consistent with much data on synapse-specific plasticity (Larsen and Sjöström, 2015), but extends this to show that synapse-specificity is present even when multiple synapses are trained simultaneously in a sparse CA3-driven pattern. Our reasoning is that although the stimuli are delivered in a spatially ordered manner in the CA3, their contact points on the CA1 PN dendritic tree will be dispersed and interleaved (Megías et al., 2001; Romani et al., 2024). Hence ‘trained’ synapses are interspersed randomly among non-trained ones participating in the non-overlap pattern, yet only the trained synapses potentiate. This is interesting, because these are the conditions under which one would expect to see substantial heterosynaptic plasticity (Oh et al., 2015) or even dendritic plasticity (Makara et al., 2009).

Computationally, synapse-specific plasticity under conditions of sparse patterned input is one of the key assumptions of pattern learning in heteroassociative networks (Field et al., 2020; Huang et al., 2004). Our findings confirm that this assumption works in the more complex CA3-CA1 network despite dendritic nonlinearities (Tran-Van-Minh et al., 2015; Ujfalussy et al., n.d.), heterosynaptic plasticity (Frey and Morris, 1997; Oh et al., 2015) and many circuits for feedforward and feedback inhibition (Lawrence and McBain, 2003). The theoretical simplification resulting from synapse specificity has functional network implications for increased network capacity (Knoblauch et al., 2010).

Our observation of strong response to the overlap pattern is equivalent to pattern completion or generalisation from neural network theory (Rolls, 2013) and can be interpreted in three ways. First, the spatial extent of optogenetic activation might be broader than the spots we illuminated and spill over to the neighbouring non-illuminated cells. This is unlikely to be a complete explanation because the images of illuminated patterns show well-defined and spatially separated spots (Figure 1B). Second, the plasticity in the three overlapping spots (out of five) may be strong enough to elicit a comparable LTP readout in the overlap pattern. We cannot resolve this possibility due to the response heterogeneity and variability. A third possibility is recurrent feedback in CA3 leading to pattern completion (Bennett et al., 1997). While we cannot rule this out, it seems unlikely because the LFP change in CA3 (Supplementary Figure 1) is small compared to the EPSP change (Figure 2, 3) (Supplementary Figure 3). We discount pattern completion in CA1 because CA1 PNs have very low recurrent connectivity (Knowles and Schwartzkroin, 1981). Overall, we interpret the strong overlap response as mostly due to the 60% of inputs which were common with the trained set.

### Heterogeneity and degeneracy in plasticity and summation

A key result from our work is the finding of substantial heterogeneity in plasticity outcomes (Figure 3, 4, 5). This was unexpected, as our stimulus protocol was designed to be a strong one with five repeats of 1 second of 100 Hz pairings of pre-vs-post activation of synapses. For comparison, three one-second bursts of 100 Hz stimuli are usually effective in eliciting strong LTP when delivered through electrical stimulation of Shaffer collaterals (Nguyen et al., 1994), and typical STDP protocols utilize ∼100 repeats (Wittenberg and Wang, 2006). Similarly, single-synapse protocols using glutamate uncaging in the hippocampal CA1 typically achieve LTP within 4 repeats followed by a plateau potential (Jain et al., 2024). Notably, all these protocols trigger LTP with high consistency.

Is the heterogeneity the outcome of differences in resting potential and background EPSP rates? While these are well correlated with learning (Figure 3G, H), we feel they are unlikely to be causative. Our LTP protocol involved elevated potentials (0 mV) and synaptic input rates (100 Hz) far beyond these small baseline physiology differences. We also do not expect that the learning outcomes are a binary consequence of different CA1 PN subtypes classified by CAMKII isoforms (Sarkar et al., 2021), since the plasticity levels were along a continuum, not bimodal (Figure 2 E). Instead we propose two possible models consistent with our observations. The first is that the learning state may be modulated at a cell-wide level due to neuromodulators (Hasselmo and Barkai, 1995), past activity, or transcriptional state (Alberini, 2009). The second is that CA1 PNs may intrinsically be heterogeneous in their physiology and connectivity, including differences in projections from CA3 and interneurons (Druckmann et al., 2014; Lee et al., 2014). Either of these might account for the small physiological differences and also the continuum of learning outcomes.

As an interesting counterpoint to the observed heterogeneity, we found that all cells exhibited the same form of sublinear summation described by divisive normalisation (Equation 2, Figure 6,(Bhatia et al., 2019)), both before and after learning. Further, most physiological properties of our recorded cells matched closely (Figure 3, methods). We suggest that CA1 PNs exhibit degeneracy with respect to their primary input-output functions, but diversity with respect to plasticity. This parallels other reports of highly similar functional outputs riding on diverse channel physiology (Marder and Prinz, 2002)

### Heterogeneity contributions to network functions

There are a number of possible ways in which heterogeneity may contribute to network properties. First, it is known that the hippocampus represents several dimensions sparsely, including space (Moser et al., 2008), time (MacDonald et al., 2011; Modi et al., 2014), and behavioral context (Nakazawa et al., 2016). This is consistent with our observation of a 40% sub-population of neurons which are more likely to participate in learning at any given time (Fig 3D). For example, place-cell tuning can be interpreted as learning a sparse input pattern of multisensory (substantially visual) input (Acharya et al., 2016), and the capacity of the network to remap using a distinct sampling of neurons relies on such sparse representations (Lian and Burkitt, 2021). Second, it is known from classical network theory that sparse representations are more effective in encoding large numbers of memories with minimal interference (Ahmad and Scheinkman, 2019). Third, heterogeneity in learning outcomes could contribute to homeostasis of overall network excitability. This happens through synaptic normalization at a single-neuron level (Mayzel and Schneidman, 2024; Turrigiano, 2008), and our findings of non-learners with enhanced post-stimulus hyperpoloarizaiton (Figure 5) suggests that heterogeneous learning could do the same across neurons.

Overall, we find that single neurons can be ‘trained’ to selectively recognize and integrate sparse input patterns, yet other neurons in the same network balance this with increased hyperpolarizing responses. We speculate that heterogeneity is a functional and dynamic attribute of plasticity in vivo, with different neurons being recruited to participate in network learning depending on slow changes in connectivity and intrinsic properties, and more rapid neuromodulation.

## Materials and Methods

### Key resources table

| Reagent type (species) or resource | Designation | Source or reference | Identifiers | Additional information |
| --- | --- | --- | --- | --- |
| Genetic reagent (M. musculus) | C57BL/6-Tg(Grik4-cre)G32-4 Stl/J | Jackson Laboratory | Stock #:006474 | Dr. Susumu Tonegawa's laboratory, MIT |
| Strain, strain background (Adeno-associated virus) | AAV9-flex-rev-ChR2-tdTomato | Adgene | Plasmid ID: 18917 | Scott Sternson lab, Janelia |

### Animals

Animal ethics clearance details:

Animal ethics Approval IDs:

NCBS-IAE-2019/16(R1): NCBS-IAE-2016/ 21 (ME); NCBS-IAE-2016/ 7 (M)

All protocols proposed have been approved by the NCBS Institutional Animal Ethics Committee. All virus lines have been approved by the NCBS Biosafety Committee.

### Animal details

We used the mouse line C57BL/6-Tg(Grik4-cre)G32-4Stl/J mice along with a Lox-P-based genetic combination that allows us to express proteins limited to CA3 pyramidal neurons. The animals were hosted in a temperature controlled specific pathogen safe environment, with a 14hr light: 10hr dark cycle, with ad libitum food and water. For obtaining the desired transgenic mouse, we crossed a transgenic male with a wild type female.

### Virus injections

The transgenic animals obtained from the crosses were injected with a virus Lox-ChR2 construct at the age of three to four weeks. The Lox construct with AAV9-flex-rev-ChR2-tdTomato (Addgene plasmid no. 18917) limits the expression of the ChR2-tdTomato to CA3 pyramidal neurons with a Grik4-Cre Transgenic mouse. The injections were performed using a sharp glass electrode inserted at the coordinates: −2.0 mm RC, +/- 1.9 mm ML, −1.5 mm DV. The volume of injection was controlled using a Picospritzer-(Picospritzer-III (Parker-Hannifin, Cleveland, OH)). By controlling the pulse width and pressure of the injections, we injected ∼500-600 nl of the virus solution with a titer of ∼5×10^12 GC/ml. We confirmed the expression of ChR2-tdTomato with epifluorescence imaging on the slices used for experiments. Post injection, animals were reared for 5-7 weeks before slice preparation.

### Slice preparation

The slices were prepared from 8-12-week-old animals (5-7 weeks post injection). Animals were decapitated after cervical dislocation, post anaesthesia with Halothane. 400 um thick slices were made from the dissected hippocampus on a vibratome (LEICA VT1200). The dissections and slicing were performed in cutting solution, artificial cerebrospinal fluid (aCSF) with (in mM): NaCl-87, KCl-2.5, NaHCO3-25, HaH2PO4.H2O-1.25, Sucrose-75, MgCl2-7, CaCl2--0.5. Once sliced, the slices were transferred to a holding aCSF with (in mM): NaCl-124, KCl-2.7, NaHCO3-26, HaH2PO4.H2O-1.25, Glucose-10, MgCl2-1.3, CaCl2-2. Both solutions were saturated with 95% O2 and 5% CO2. After an incubation of 45 mins, Slices were transferred to the recording chamber with holding-aCSF perfusion at 32+/- 1.5°C.

### Electrophysiology

We used an infrared differential interference contrast (DIC) microscope(Olympus BX51Wi) with 40x (Olympus LUMPLFLN, 40XW) water immersion objectives to visualise the cells. The expression of ChR2-tdTomato was confirmed with the fluorescence imaging system on the same scope. The electrophysiology data was acquired using Axon™ Digidata® 1440B digitizer and MultiClamp 700B amplifier (Molecular Devices, Axon instruments). We used 20KHz sampling frequency for pattern frames and 10KHz for single-point frames. A Bessel bandpass filter between 10-10KHz was used for the acquisition of electrical signals. We performed whole-cell patch recordings from CA1 pyramidal cells and single electrode field recordings from the CA3 region. The patch electrodes are made from thick-walled borosilicate glass capillaries(OD-1.56, ID-0.86mm, Sutter instruments). The resistance range of all recording electrodes was between 2.8-5 MOhms. We used a P-1000 pipette puller from Sutter Instruments, Novato, CA, with a 2.5mm and 3mm box filament for electrode pulling.

Electrodes were filled with (mM): K-Gluconate-130, NaCl-5, HEPES-10, Na4-EGTA-1, MgCl2-2, Mg-ATP-2, Na-GTP-0.5, Phosphocreatine-10, pH adjusted to 7.3 and osmolarity adjusted to ∼290 mOsm. CA1 pyramidal neurons were maintained close to −65 mV, and a small current injection of <20 pA was delivered to reach this range in the current clamp protocols. Cells were considered for analysis based on the stability of series-resistance observed. We set the following inclusion criteria for recordings: input resistance change remained below 15% from the start), and membrane voltage stayed within 1.5 mV from the median resting membrane potential.

To induce plasticity, we paired an optical stimulation at CA3 PNs with a 1nA, 2ms current injection at CA1. The optical stimuli led the current injection by 5ms. This was presented via 100 pulses at 100Hz and repeated five times with a 0.22 s separation.

### Optical stimulation setup

To stimulate the ChR2 expressing cells, we used a Digital Micromirror Device (DMD) clubbed with an LED (Polygon400 (DMD projector) and BioLED Light Source Control Module (Mightex, Canada). The assembly was installed in the light path of the microscope and controlled with an external trigger generated from the amplifier. The setup was equipped with 470 nm and 590 nm wavelength LEDs. We used 470 nm for all our experiments. The DMD pattern changing rate was from 0 Hz to 4 KHz for binary images. This is sufficient to generate any frequency of optical stimulation we would like to use in our protocols. The DMD had a resolution of 684 x 608 pixels/points; this translates to 0.49 *µ*mx0.32 *µ*m per pixel size under 40x magnification. Thus, the resolution is substantially finer than single neuron soma size (∼20 *µ*m). The LED brightness can be controlled from 0-100% using the LED module.

We used 100% brightness for all our stimuli. A photodiode connected to an amplifier was installed in the turret to validate the projection state from Polygon. We utilised the proprietary software supplied with Polygon to project patterns and waveforms needed for optical stimulation.

### Patterned optical stimulation

Once the CA1 cell was patched, we moved the objective field of view from CA1 to the CA3 region for optical stimulation. Each projection of a frame was 2 ms long to elicit a spiking response in CA3 pyramidal neurons. The frames with patterns consisted of five bright points in a 24×24 grid size, and each grid point was calculated to span 13 x 7 *µ*m under a 40x objective. The stimulus pattern was a linear array of illumination spots positioned to cover the CA3 cell body region. There was a 13um separation between adjacent points, and each frame was presented with a 5s time gap. There were a total of three patterns, with one used for training, one with 60% overlap with the trained pattern and one with no overlap with the trained pattern. Our stimulus sequence included first the three individual patterns,(training, overlap and non-overlap) with five points each. Then we delivered each of the twelve points one at a time. For all three patterns we performed three repeats in one block and the same was done for the 12 points.Successive frames were presented at 0.2 Hz. From a field electrode placed in the CA3 region, we measured the field response elicited by the optical stimulation.

To measure the post training responses of CA1 PN to patterns and points presented at CA3, we waited for the CA1 PN resting membrane potential to come back to the pre-training value. The time point at which it came back to pre-training resting membrane potential was considered as 0 minutes and from there on, the measurements were done at 10 minutes gap.

### Analysis pipeline

The electrophysiology recording data files saved from the amplifier and digitizer were accessed using “neo”, an open source library in python. The analysis was performed using custom scripts written in python. The script is available in the repository (github link:https://github.com/anzalks/single_neuron_pattern_learning_paper).

Bootstrap analysis between γ-pre and γ-post was performed by pooling the raw pre and post datasets in the form of individual trial coordinates for (expected sum, observed sum). N = len(dataset) samples were taken with replacement from this pooled dataset, and used to calculate a sample each for γ-pre and γ-post. N was in the range 24 to 30. These values were recomputed 10,000 times and in each case their difference was computed to form the null-hypothesis distribution for γ difference. The fraction of this distribution that exceeded abs(γ-pre - γ-post) was taken as the p-value. In each case γ was computed by nonlinear least-squares fitting of the respective datasets to equation 2 using the routine scipy.optimize.curve_fit() from scipy version 1.14.1.

## Supplementary Data and Figures

### Comparison of different EPSP trace properties over time points

**Supplementary Figure 1.**
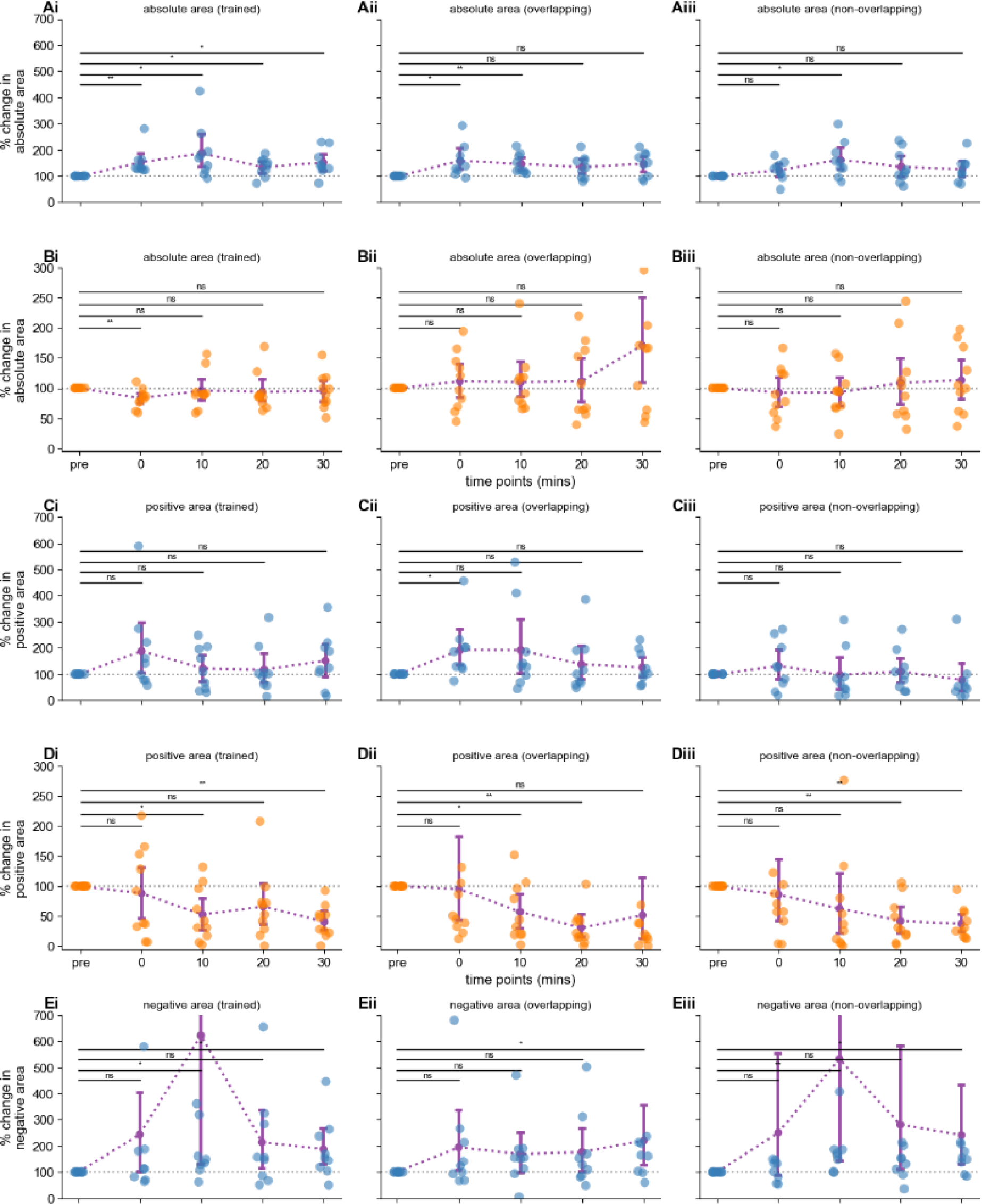
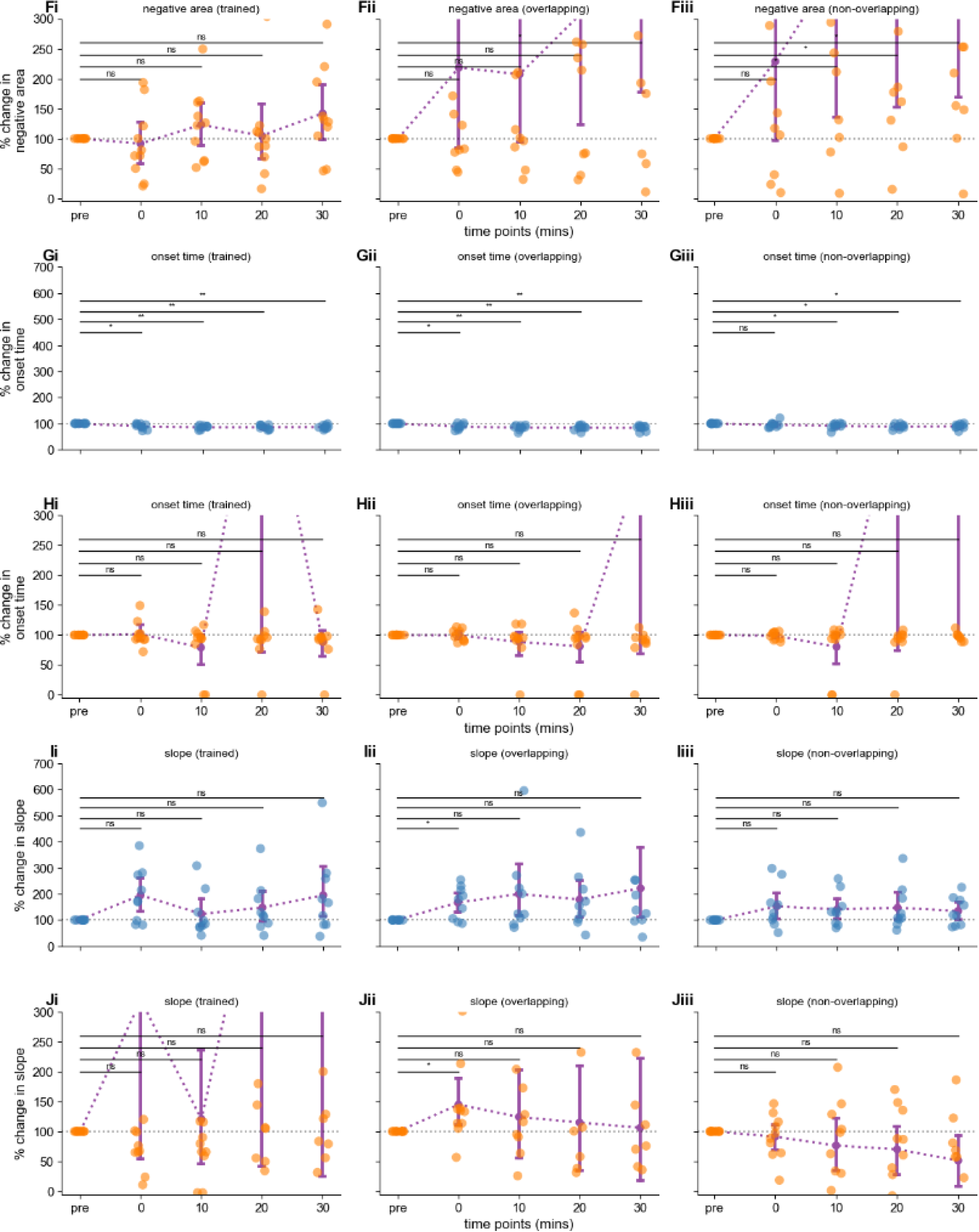
Percent change in various EPSP readouts in response to training across multiple time points for learners and non-learners. We showed that multiple readouts of EPSP have comparable effects in response to LTP induction. In each case we estimated the readout for multiple time points (pre, 0 min, 10 min, 20 min, and 30 min) following the training protocol, for and for learners (blue) and non-learners (orange). Each parameter was assessed under three conditions: trained (left panels), overlapping (middle panels), and non-overlapping (right panels). The significance of differences between time points was assessed using the Wilcoxon signed-rank test. Individual neurons are shown as colored dots, with error bars denoting the mean ± SEM.

- **(A)**: Absolute area under the curve for learners.

- **Ai (Trained)**: At 30 mins, learners (blue) showed a significant sustained increase compared to pre-training (p = 0.012 at 30 min).
- **Aii (Overlapping)** Overlapping pattern elicited rapid enhancement in response immediately after training(p= 0.012), but no sustained increase at 30 mins.
- **Aiii (Non-overlapping)**: No significant changes at either 0 min or 30 min for both groups.
- **(Bi)**: Similar to A, for non-learners.

- **Bi (Trained)**: At 0 min, non-learners showed a significant decrease (p = 0.01), indicating immediate depression. By 30 min, there were no significant differences, indicating a return to baseline.
- **Bii (Overlapping)** & **Biii (Non-overlapping)**: No significant changes at either time point for either group.
- **(Ci, Cii, Ciii): Positive area learners**

- **Ci (Trained)**: At 0 min, learners show no significant increase. By 30 min, the positive area normalizes, with no significant difference from pre-training. Non-learners show no significant changes at either time point.
- **Cii (Overlapping)**: at 0 min there is an increase(p=0.012).No significant changes at 30 min.
- **Ciii (Non-overlapping)**: No significant changes at 0 min to 30 min.
- **(Di, Dii, Diii): Positive area non-learners**

- All patterns except overlapping patterns show a decrease in positive area over time.
- **(Ei, Eii, Eiii): Negative area learners**

- Negative area increased over time for all patterns.
- **(Fi, Fii, Fiii): Negative area non-learners**

- Except for the trained pattern the negative area increased significantly over time(p=0.037 and 0.014 for overlapping and non-overlapping patterns respectively).
- **(Gi, Gii, Giii): Onset time (time to reach 0.5 mV post stimulus) learners**

- For all the patterns, the onset time is reduced over time.
- **(Hi, Hii, Hiii): Onset time (time to reach 0.5 mV post stimulus) non-learners**

- For all the patterns, the onset time did not show any significant difference over time.
- **(Ii, Iii, Iiii): Slope learners**

- No significant change in slope over time except for the 0 min time point in overlapping pattern(p=0.049)
- **(Ji, Jii, Jiii): Slope non-learners**

- No significant change in slope over time except for the 0 min time point in overlapping pattern(p=0.049)

### CA3 field responses increase following LTP induction, but not enough to change EPSP LTP outcomes

**Supplementary Figure 2.**
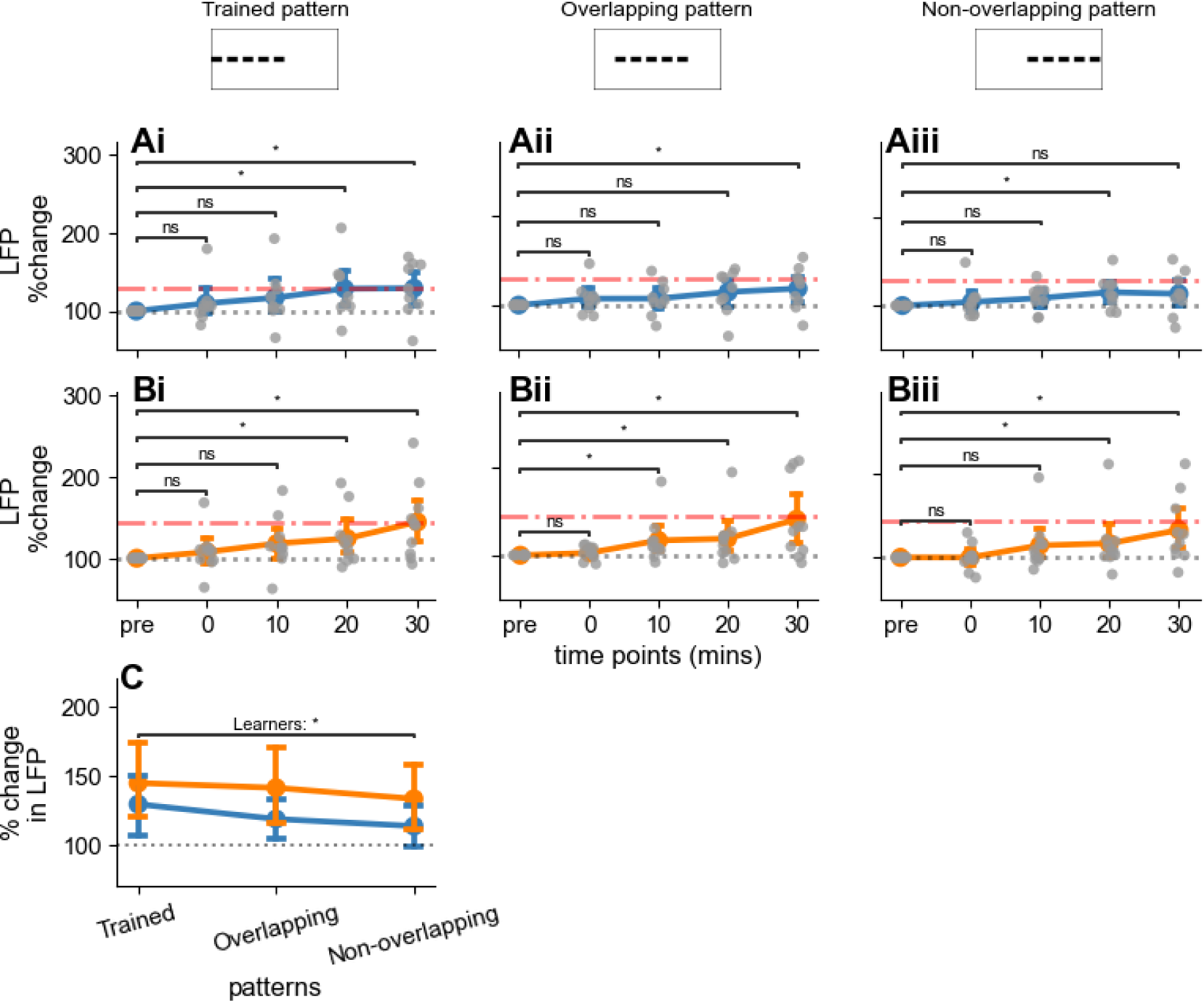
Field response changes in CA3 neurons following LTP-induction. - **(A)** Percentage change in field response over time for learners across the three patterns. Responses are normalized to baseline. Trained (Ai, p = 0.04) and overlap (Aii, p = 0.039) patterns elicit a significant elevation at the 30 minute time point. Non overlapping patterns do not (Aiii, p = ns).
- **(B)** Same as A, for non-learnerns. All three patterns elicit a significant potentiation in LTP, showing that the difference between learners and non-learners is not due to the stimulus. Bi: Trained, p = 0.013, Bii: Overlap, p = 0.13, Biii: non-overlap, p = 0.19
- **(C)** Summary of percentage change in field response at 30 minutes post-training across patterns for both learners (blue) and non-learners (orange). The bar plot shows the mean percentage change ± SEM. learners show a significant increase in LFP amplitude between the trained pattern and non-overlapping patterns(p=0.04). Wilcoxon signed rank test was used in all cases.

### Pattern selectivity and cell classification holds even with EPSP amplitudes normalised to LFP amplitudes

**Supplementary Figure 3.**
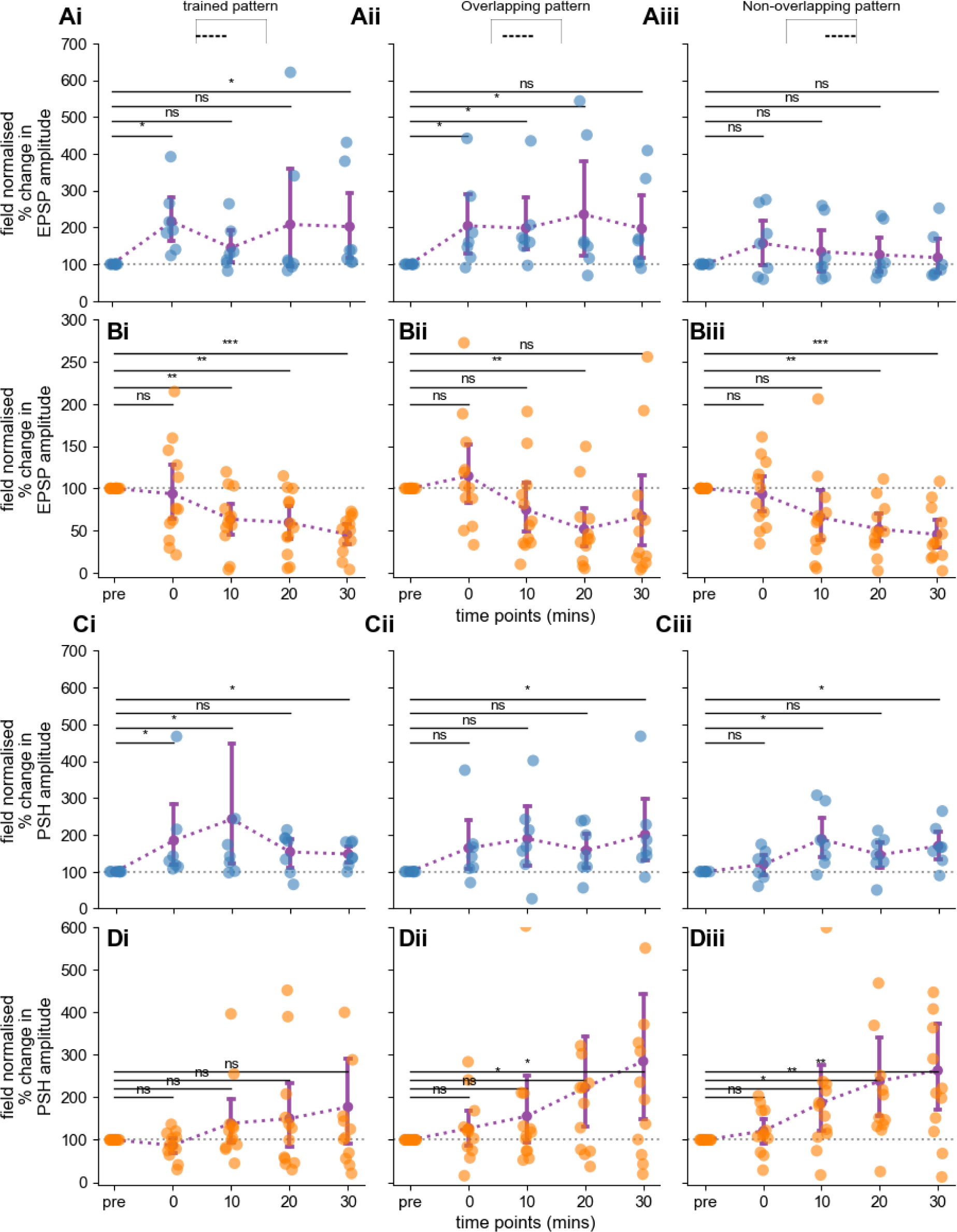
LFP normalised EPSP amplitude and PSH changes following optogenetic stimulation patterns for learners and non-learners. **(Ai-Aiii)** Quantification of change in EPSP amplitude in learner cells (n=6). Amplitudes reported as percentage change normalized to baseline for trained (Ai), overlapping (Aii), and non-overlapping (Aiii) stimulation patterns across time points. Individual points represent single cells, lines and error bars indicate the mean ± SEM. at 30 minutes post training, the trained ( p=0.015) pattern has a significant increase. Overlapping (p=ns) and non-overlap (p=ns) patterns have no significant change by the Wilcoxon signed rank test. **(Bi-Biii)** Same as A for non-learners (orange, n=12). Trained (p=0.0004) and non-overlap (p=0.0009) patterns show a significant decrease, Wilcoxon signed rank test. **(C)** Quantification of the percentage change from baseline in PSH amplitude for the trained **(Bi)**, overlapping **(Bii)**, and non-overlapping **(Biii)** stimulation patterns across time points (pre-training, 0, 10, 20, and 30 minutes post-training) in learner cells (blue, n=6). Each point represents an individual cell, and lines with error bars indicate the mean ± SEM. Trained (p=0.03), overlap (p=0.03) and non-overlap (p=0.03) patterns show a significant increase (Wilcoxon signed rank test). The purple dashed lines represent the trend of PSH amplitude changes over time. **(D)** Similar to C, for non-learners.overlapping (p= 0.027) and non-overlap (p=0.0068) patterns show a significant decrease, Wilcoxon signed rank test.

### Synaptic summation becomes more linear for trained patterns after LFP normalisation

**Figure 4.**
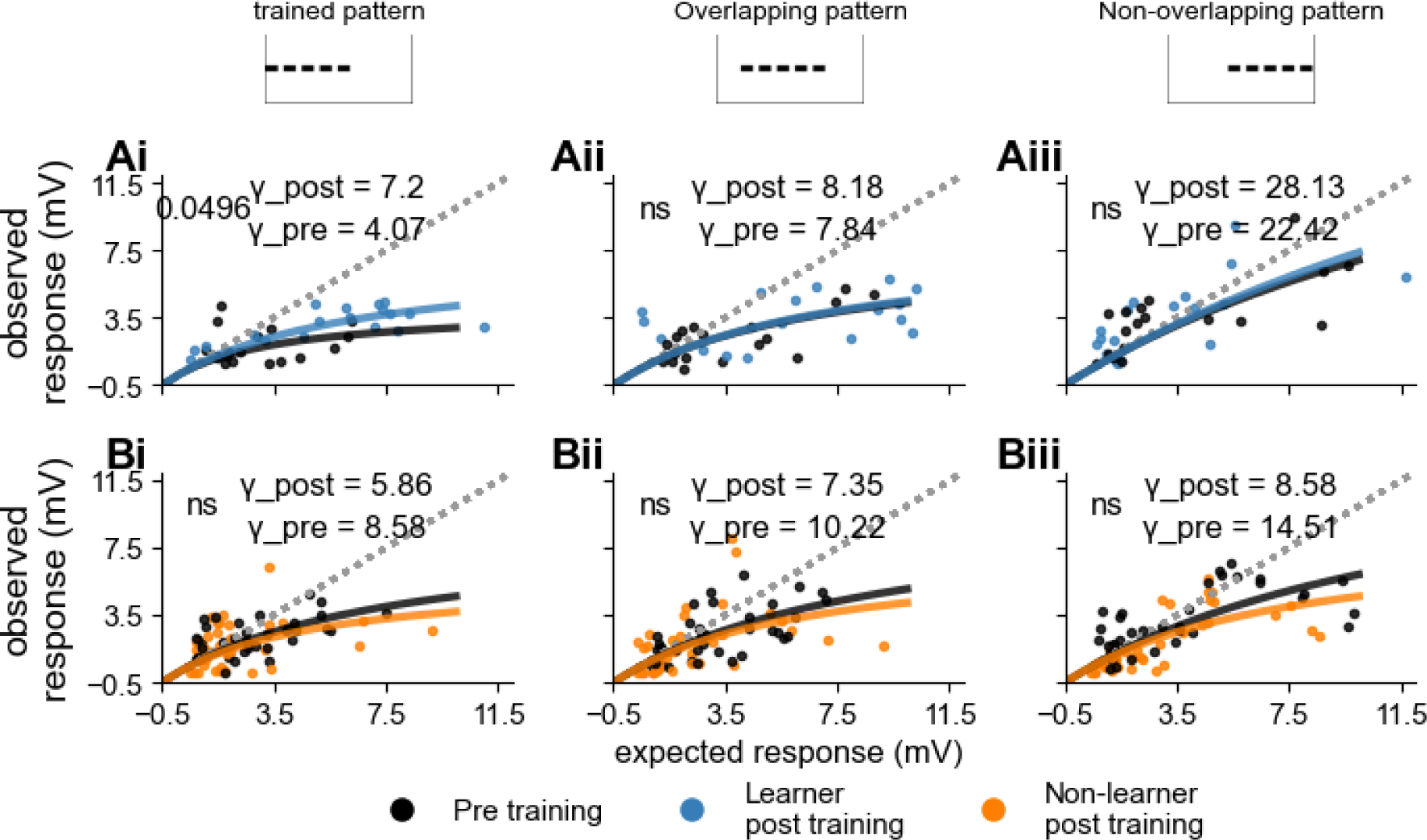
Sublinear Summation and Divisive Normalization in CA1 Neurons Across Different Patterns after normalising EPSPs with LFP. **(A)** Results from learners(n=6, 3 trials per pattern), with fits for γ-pre (black) and γ-post (blue). A linear sum is represented by the dotted line. **(Ai)**For trained pattern (γ-pre = 4.07, γ-post = 7.2, p=0.047) showed a just significant increase in gamma post training **(Aii)** Overlapping pattern (γ-pre = 7.84, γ-post = 8.18, p=ns) and **(Aiii)** Non-overlapping pattern (γ-pre = 22.42, γ-post =28.13, p=ns) have no significant change in the summation. **(B)** Results from non-learners (n=12, 3 trials per pattern).**(Bi)** Trained pattern (γ-pre = 5.86, γ-post = 8.58, p=ns). **(Bii)** Overlapping pattern (γ-pre = 7.35, γ-post = 10.22, p=ns) and **(Biii)** non-overlapping pattern (γ-pre = 14.51, γ-post = 8.58, p=ns) γ does not change significantly. We also did not see any significant difference between the gammas of trained and non-overlapping patterns.

### Synaptic responses and intrinsic properties of CA1 neurons after classification based on LFP normalised EPSP amplitudes

**Supplementary Figure 5.**
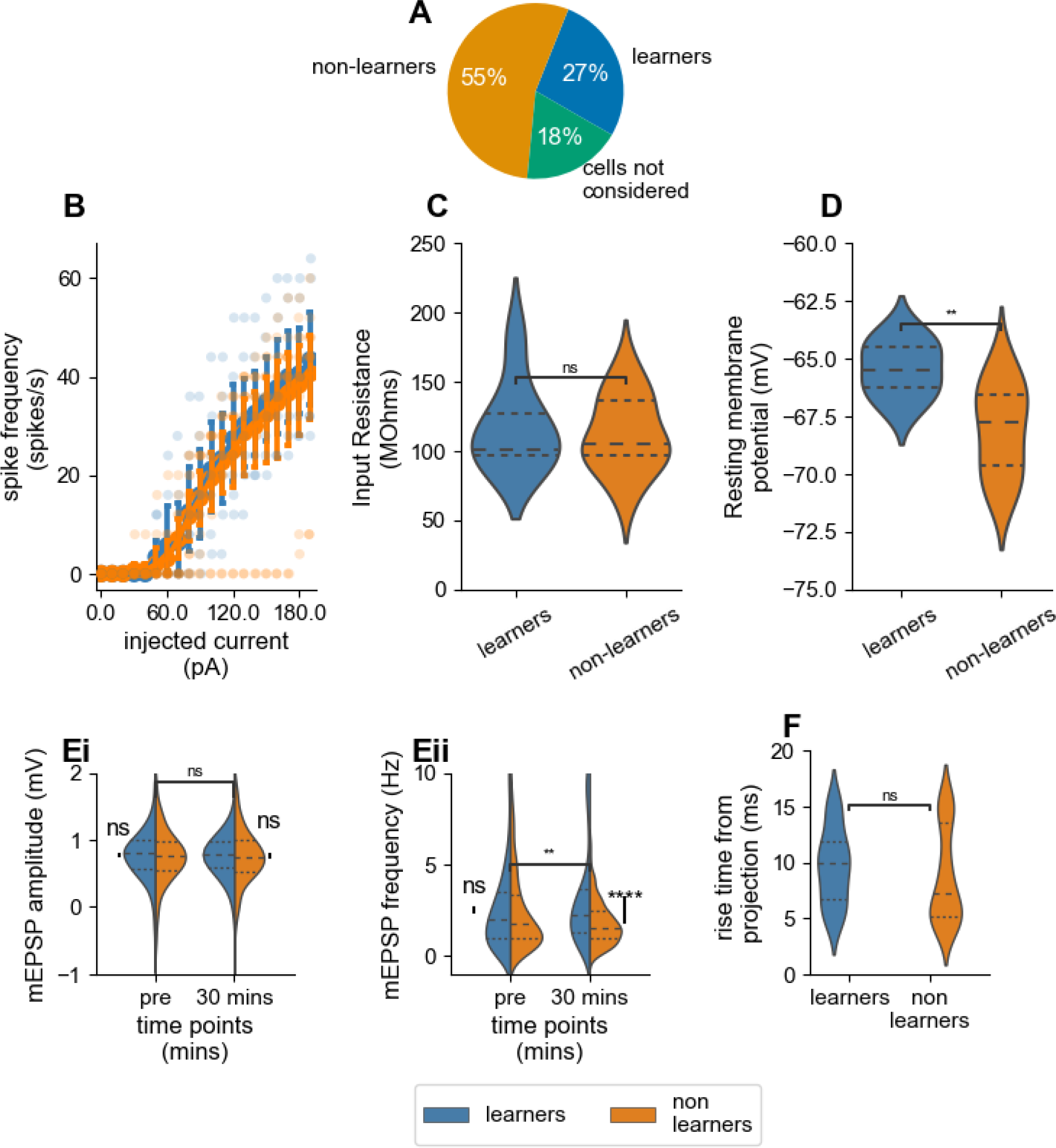
Changes in synaptic responses and intrinsic properties of CA1 neurons during pattern learning based on LFP normalised EPSPs. **(A)** Pie chart summarizing the distribution of cells classified as learners (∼27%, n=6), and non-learners (∼55%, n=12). Four cells (∼18%) were excluded because their EPSP amplitudes were less than 0.5 mV, or because they spiked after training. **(B)** Input-output relationship between spike frequency and injected current for learners (blue) and non-learners (orange). The curves did not differ. Error bars represent mean ± SEM. **(C)** Comparison of input resistance (MΩ) between learners and non-learners. No significant differences were observed (Mann-Whitney U test). **(D)** Comparison of resting membrane potential between learners and non-learners. Learners exhibited a significantly more depolarized resting membrane potential compared to non-learners (p = 0.007,Mann-Whitney U test). **(E)** Miniature EPSP (mEPSP) analysis in learners and non-learners, compared before and 30 minutes after training.**(Ei)** mEPSP amplitude comparison shows no significant differences. **(Eii)** Comparison of mEPSP frequency shows a significant difference between the learners and non learners (p<0.0001, Mann-Whitney U test). The pre and post-training frequency shows an increase in learners(p=0.009). **(F).** Response rise time, measured as time from delivery of optical stimulus till reaching 0.25 mV depolarization. Learners and non-learners did not differ (Mann-Whitney U test).

### Non-learners show an increase in LFP amplitudes over time, once cells are classified based on LFP normalised EPSPs

**Supplementary Figure 6.**
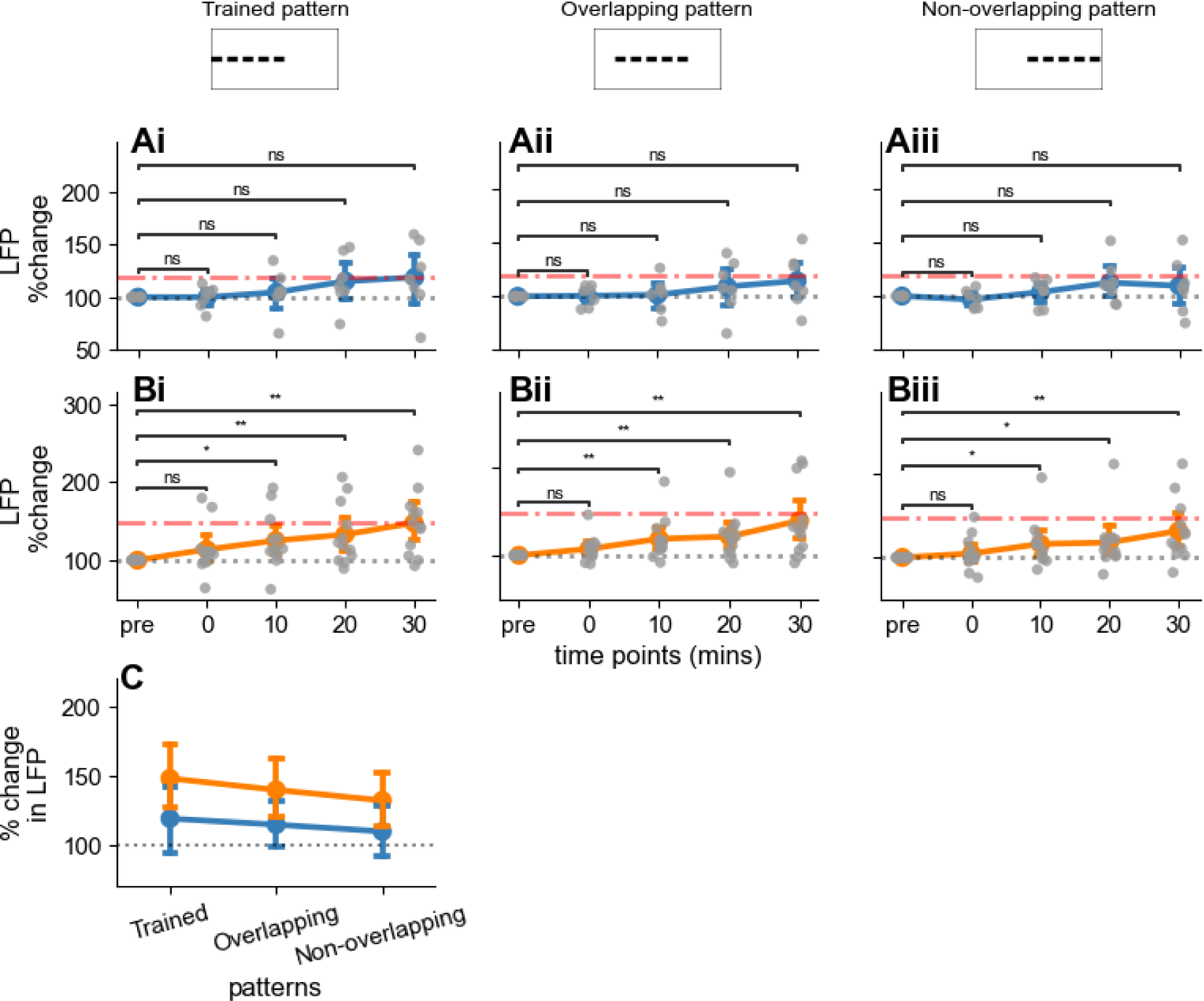
Field response changes in non-learner’s CA3 neurons following LTP-induction. **(A)** Percentage change in field response over time for learners across the three patterns. Responses are normalized to baseline(n=6). Trained, overlapping and non-overlapping patterns do not show any significant change in the LFP amplitude over time. **(B)** Same as A, for non-learnerns. All three patterns elicit a significant potentiation in LTP, showing that the difference between learners and non-learners is not due to the stimulus. **(Bi)** Trained, p = 0.0034, **(Bii)** Overlapping, p = 0.0034, **(Biii)** non-overlapping, p = 0.007( Wilcoxon signed rank test) showed significant increase at 30 minutes post training. **(C)** Summary of percentage change in field response at 30 minutes post-training across patterns for both learners (blue) and non-learners (orange). The bar plot shows the mean percentage change ± SEM. The patterns did not show any significant difference in LFP amplitude between the trained pattern and non-overlapping patterns(p=ns). Wilcoxon signed rank test was used in all cases.

### Quantification of desensitisation of ChR2 and Na+ channel inactivation during training

**Supplementary Figure 6.**
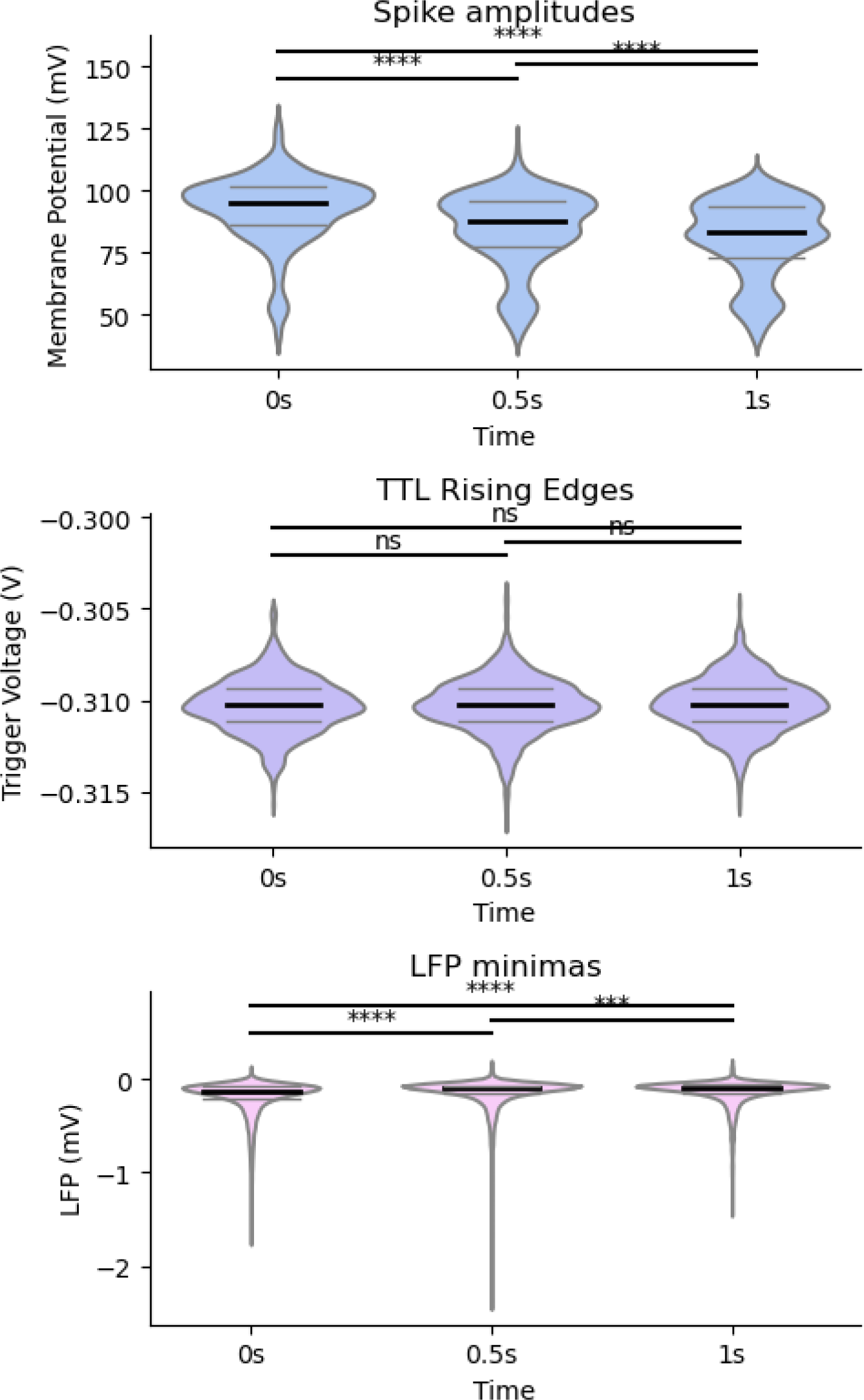
Violin plots of spike amplitudes, TTL rising edges, and LFP peaks at different time points. Each panel shows the distribution of values for three time points: 0s, 0.5s, and 1s. Each plot includes pairwise Mann-Whitney U test results displayed above the violin plots, with lines connecting the compared groups. The p-values are shown alongside the significance markers.

- **Top Panel (Spike amplitudes)**: Membrane potential values from recorded spikes. The black horizontal lines indicate the median, and the grey lines represent the 25th and 75th percentiles. Spike amplitudes showed a significant decline over time.
- **Middle Panel (TTL Rising Edges)**: Trigger voltage values at different time points. The central lines indicate median values with interquartile ranges. Pairwise comparisons between time points show no significant differences, as denoted by “ns” labels.
- **Bottom Panel (LFP Peaks)**: Local field potential (LFP) peak measurements. Median and interquartile ranges are represented similarly. The comparison between the 0s – 0.5, 0.5 – 1s and 0s – 1s time point comparisons shows a highly significant reduction in the LFP amplitude.

## Acknowledgements

NCBS-TIFR receives the support of the Department of Atomic Energy, Government of India, under Project Identification No. RTI 4006. The project also had support from SERB Core Research Grant (CRG), Govt of India, file no. CRG/2022/003135. We thank Arvind Kumar, Rishikesh Narayanan and Sufyan Ashhad for general suggestions over the years, Gilad Silberberg, Sho Yagishita in helping with standardising the field electrode measurements and the plasticity induction protocol, and Anchal Bhatia, Sahil Moza and Anal Kumar for the discussions to formulate the project and set up the analysis pipeline in the early stages of the project.

## Author contributions

AKS performed the experiments, did the analysis, and wrote the paper. USB supervised the project, did the analysis, and wrote the paper. USB and AKS obtained the funding.

